# Construction of Complete Telomere-to-Telomere Genome Assembly of the Rabbit Using Haploid Embryonic Stem Cells

**DOI:** 10.1101/2025.03.26.645123

**Authors:** Yunga A, Junjie Mao, Shengyu Hu, Manya Yu, Yuanxi Yang, Yawei Zhang, Tong Qiu, Longquan Quan, Hongqiang Zhao, Quanjun Zhang, Kang Yu, Yinghua Ye, Haizhao Yan, Nan Li, Xiong Yang, Xiawen Yang, Rufan Zhang, Guangyao Zhai, Minxian Wang, Liangxue Lai

## Abstract

The rabbit owns commercial importance in meat and fur production, and also has long served as a valuable animal model in biomedical research. Yet, the complete assembly of a high-quality rabbit reference genome has not been established. Here, we present a telomere-to-telomere (T2T) genome assembly of the New Zealand White (NZW) rabbit, the most complete and accurate rabbit genome to date. Using a haploid embryonic stem cells (haESCs) to overcame the challenges of homologous sequence interference and structural variation, we generate a genome with a contig N50 of 137.71 Mb, anchoring all chromosomes with only three gaps, through integrating long-read sequencing (PacBio HiFi, Oxford Nanopore), Hi-C, and Illumina data. Quality assessments reveal near-complete coverage of the BUSCO mammalian gene set, with 99.61% completeness. Notably, this complete assembly allowed for the first comprehensive annotation of the rabbit immunoglobulin loci and major histocompatibility complex region, which were previously unresolved in earlier genome versions. The NZW-T2T assembly based on haESCs not only provides a critical resource for advancing rabbit-based research, but also set a new benchmark for de novo genome assembly for other animals with complex genome.

## Introduction

The European rabbit (Oryctolagus cuniculus), including its domesticated breeds, is distributed across more than 70 countries (Carneiro et al., 2011). Due to its intermediate body size, physiological similarities to humans, justifiable expenditure, and commercial importance in meat and fur production, the rabbit serves as an ideal animal model for cardiovascular, ophthalmological, toxicological, immunological, and infectious studies. For example, rabbits can be used as a preferred model for non-infectious diseases such as atherosclerosis, intestinal immunity, reproduction, lupus, arthritis, cancer, genetic cardiac channelopathies, Alzheimer’s disease, and so on (Esteves et al., 2018), as well as for infectious diseases such as Mycobacterium tuberculosis (Subbian et al., 2015), syphilis (Lithgow et al., 2017), and HIV-1 (Speck et al., 1998; Yamamura et al., 1991). Particularly, because the immune genes of rabbits are more similar to those of humans than those of rodents, and research on rabbit immunoglobulins has significantly advanced our knowledge of antibody structure, function, and regulation (Pinheiro et al., 2016; Soares et al., 2022). The complete assembly of a high-quality rabbit reference genome would greatly facilitate above ongoing research. However, the complete assembly of a high-quality rabbit reference genome is not available as far.

The first assembly of the rabbit genome, OryCun1.0, was constructed using 2x coverage sequencing, which included ∼1.8x coverage from 4 kb paired-end reads and ∼0.2x from 40 kb Fosmids (Lindblad-Toh et al., 2011). Due to the technological constraints of sequencing and assembly methods at that time, OryCun1.0 was of relatively low quality, with a total genome length of only 2.08 Gb and an N50 contig size of 3.3 kb. Later, based on OryCun1.0, OryCun2.0 was generated with additional paired-end reads, containing an N50 contig size of 64.7 kb and a length of 2.18 Gb sequence in anchored scaffolds (Carneiro et al., 2014). More recently, with the advent of third-generation sequencing technologies, the UM_NZW_1.0 assembly was generated based on long-read data from the New Zealand White (NZW) rabbit (Bai et al., 2021). While this assembly greatly improved the continuity and completeness of the rabbit genome (compared to OryCun2.0) and maintained high accuracy, it still has 21,334 contigs, with 562.64 Mb of sequences unanchored to chromosomes, indicating significant discontinuity.

The telomere-to-telomere (T2T) assembly of a human genome (Nurk et al., 2022), and other model organisms like mice (Liu et al., 2024) and important crops like maize (J. Chen et al., 2023), providing unprecedented insights into complex genomic regions through complete or near-complete assemblies. However, construction of the telomere-to-telomere (T2T) assembly is a complex procedure because they used diploid or polyploid cells for sequencing, which may cause the mutual interference between homologous sequences, segmental duplications, and sequencing errors (Li & Durbin, 2024). The supplementary data, such as Hi-C contact matrices or allelic information from parents, can help separating haplotypes in some extent, current genome assemblers were unable to separate them reliably (Koren et al., 2018). Haploidy with a single set of chromosomes, presents a potential solution to this challenge. The first T2T human genome assembly was achieved by utilizing the complete hydatidiform mole cell line (CHM13), known for its near-uniform homozygosity across the genome (Nurk et al., 2022). However, certain heterogeneities still exist in hydatidiform mole cells, such as a megabase-scale heterozygous deletion within the rDNA array on chromosome 15, introducing variability in the specific regions and complicating assembling process (Rautiainen et al., 2023). The haploid embryonic stem cells (haESCs) with only one set of chromosomes containing complete sequence of each chromosome, which have been successfully established in several species such as fish (Yi et al., 2009), mice, rats, and humans (Wang et al., 2024), representing true haploidy. The use of haESCs can overcome several challenges from switch errors, a mixture of haplotypes of large-scale or complex structure variants, and sequencing errors, is expected to greatly enhance the accuracy and efficiency of genome assembling (Nurk et al., 2022). Recent pioneering research on the mouse T2T genome has demonstrated that haploid embryonic stem cells (haESCs) offer significant advantages in genome sequence assembly (Liu et al., 2024).

Here, we generated the first T2T genome assembly of New Zealand White rabbit (NZW-T2T), by using a complete haploid cell line generated from rabbit parthenogenetic blastocysts. After assembling, scaffolding, and merging, we generated the first near T2T genome assembly of New Zealand White rabbit (NZW-T2T), with all contigs anchored continuously on the chromosomes, except for chromosome 15, contains four contigs separated by three gaps, and is the most accurate and complete rabbit genome assembly to date. Based on our NZW-T2T, we rebuilt the entire rabbit immunoglobulin locus for the first time and annotated a good number of novel variable genes that are invaluable for the study of immunogenetics. Additionally, we analyzed the arrangement and characteristics of the rabbit MHC locus from an evolutionary perspective and found that rabbits may be an ideal candidate recipient for producing human cells heterologously.

## Results

### Rabbit T2T genome assembled using haESCs

In order to have a haploid genome for assembly, we derived rabbit haESCs (rbHaESCs) from rabbit parthenogenetically developed haploid blastocysts. Our results indicated that rbHaESCs contain 22 chromosomes (Fig. 1A), representing half of the total rabbit chromosome numbers, and exhibit minimal heterogeneity within the population as anticipated (Fig. S1). Since a complete genome assembly for rabbits has not yet been constructed, we estimated the size of rabbit genome through flow cytometry, using mouse haESCs as a reference due to the close evolutionary relationship between Lagomorpha and rodents (Graur et al., 1996). Our findings confirmed the haploid status of rbPhESCs, as most cells were enriched under the 1N peak, and demonstrated that the DNA content of rabbits closely resembles that of mice, as indicated by the nearly overlapping haploid and the diploid peaks between rabbit and mouse (Fig. 1B).

**Fig. 1.**
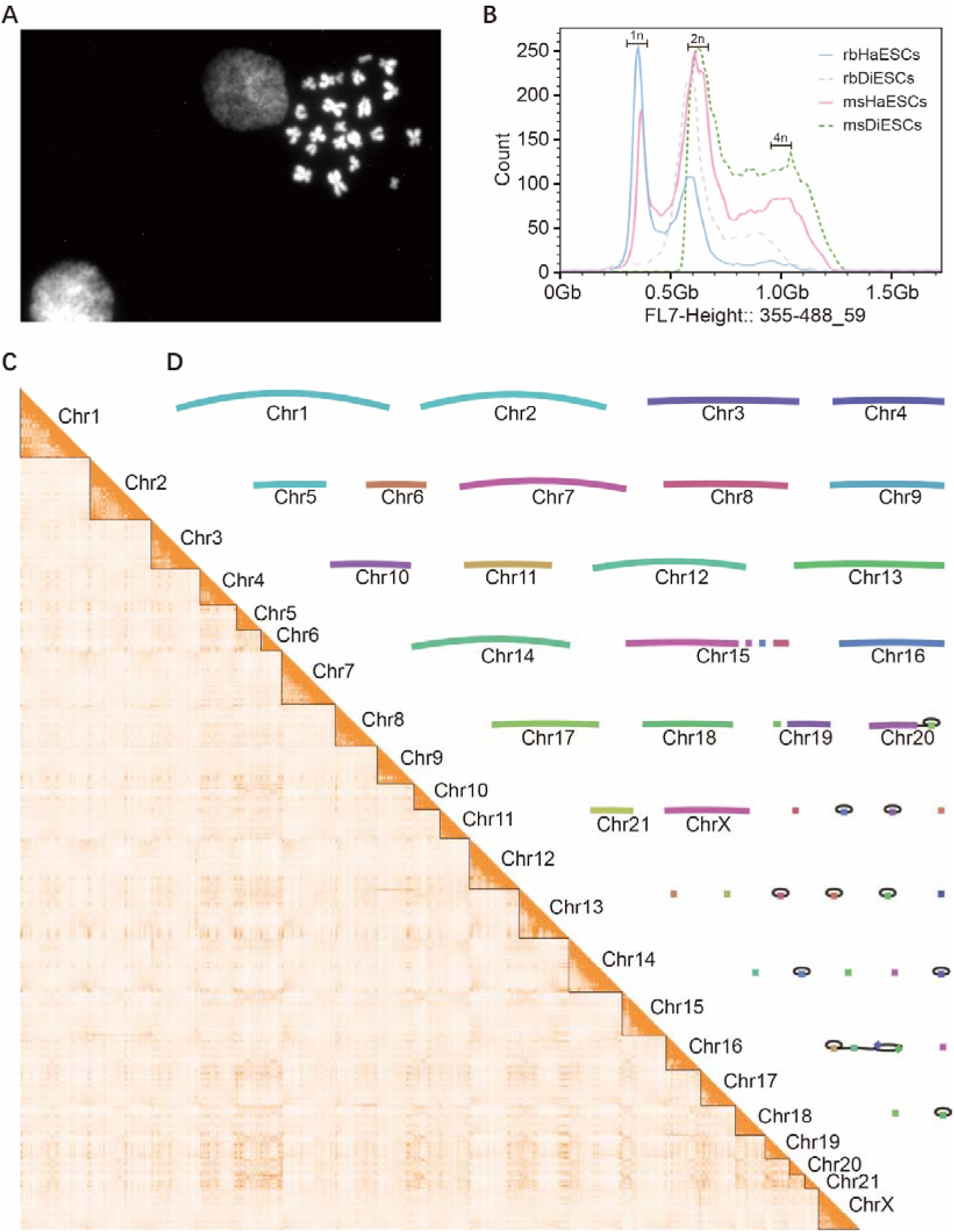
Summary of the near-complete NZW-T2T rabbit genome assembly. (**A**) Chromosome spreads of rbPhESCs, each containing 22 chromosomes, half the number found in normal rabbit somatic cells. (**B**) Genomic content analyzed by FACS, mouse haploid and diploid embryonic stem cells were used as control. The 1n and 2n chromosome sets for haploid and 2n and 4n chromosome sets for diploid ESCs are indicated at the peak positions. rbHaESCs, rabbit haploid embryonic stem cells; rbDiESCs, rabbit diploid embryonic stem cells; msHaESCs, mouse haploid embryonic stem cells; msDiESCs, mouse diploid embryonic stem cells. (**C**) Chromosomal Hi-C heatmap of the NZW-T2T genome assembly. (**D**) A Bandage visualization of the assembly graph, where nodes represent unambiguously assembled sequences scaled by length and edges correspond to the overlaps between node sequences.

We then sequenced rbHaESCs using different technologies helpful for assembly, including 150.49 Gb Oxford Nanopore ultralong-read sequencing, 139.39 Gb PacBio HiFi Revo sequencing, 257.63 Gb Hi-C sequencing, and 131.8 Gb Illumina PCR-Free whole genome sequencing (Table S1-S5). We generated two assemblies: one named asm_hia, which was constructed using hifiasm-ul (Cheng et al., 2024), the same assembler employed for the Human Pangenome Project; and the other named asm_vko, which was generated using the Verkko pipeline (Rautiainen et al., 2023). Both assemblies were then scaffolded using Hi-C contact matrices to cluster spatially proximate contigs, enabling the formation of chromosome-scale scaffolds (Fig. S3, Fig. S4). After that, 18 scaffolds in asm_hia contained telomere-specific motifs at both ends without gaps (Fig S5), indicative of complete chromosomes, compared to 12 scaffolds in asm_vko (Fig S6) (Table S6). Therefore, we used the asm_hia version as the main assembly, combining chromosomes 19 and X from asm_vko to asm_hia assembly to generate the final assembly. After polishing, we obtained the Telomere-to-Telomere assembly, with most of the chromosomes displaying a linear structure in the resulting graph (Fig. 1D). Notably, two gaps in chr19 were resolved using Oxford Nanopore Technology (ONT) reads via MaSuRCA (Zimin et al., 2013). However, for the gaps on Chr15, existing gap-filling tools failed to resolve it, resulting in three retained gaps in the final genome assembly. The Hi-C interaction heatmap showed that each chromosome of NZW-T2T displays a clear boundary, indicating a complete and high-accuracy genome assembly (Fig. 1C).

### Genome quality assessment

To evaluate assembly quality, we compared key metrics of the NZW-T2T with those of existing rabbit assemblies, including OryCun2.0, OryCun3.0, and UM_NZW_1.0. Our results demonstrated that the NZW-T2T assembly contains 9,031 complete and single-copy BUSCO mammalian genes, 159 complete and duplicated BUSCO mammalian genes, and only 20 fragmented BUSCO mammalian genes. The number of missing BUSCO mammalian genes is remarkably low, at just 16. Overall, the number of complete BUSCO mammalian genes in NZW-T2T reaches 9,190, representing 99.61% of the entire BUSCO mammalian gene set. In comparison, the corresponding percentages for OryCun2.0, CryCun3.0, and UM_NZW_1.0 are 93.04%, 99.14%, and 96.39%, respectively. Notably, NZW-T2T exhibits the fewest fragmented BUSCO mammalian genes and the lowest number of missing BUSCO mammalian genes among these assemblies. This highlights that the genome of NZW-T2T achieves an exceptionally high level of BUSCO completeness (Fig. 2A).

**Fig. 2.**
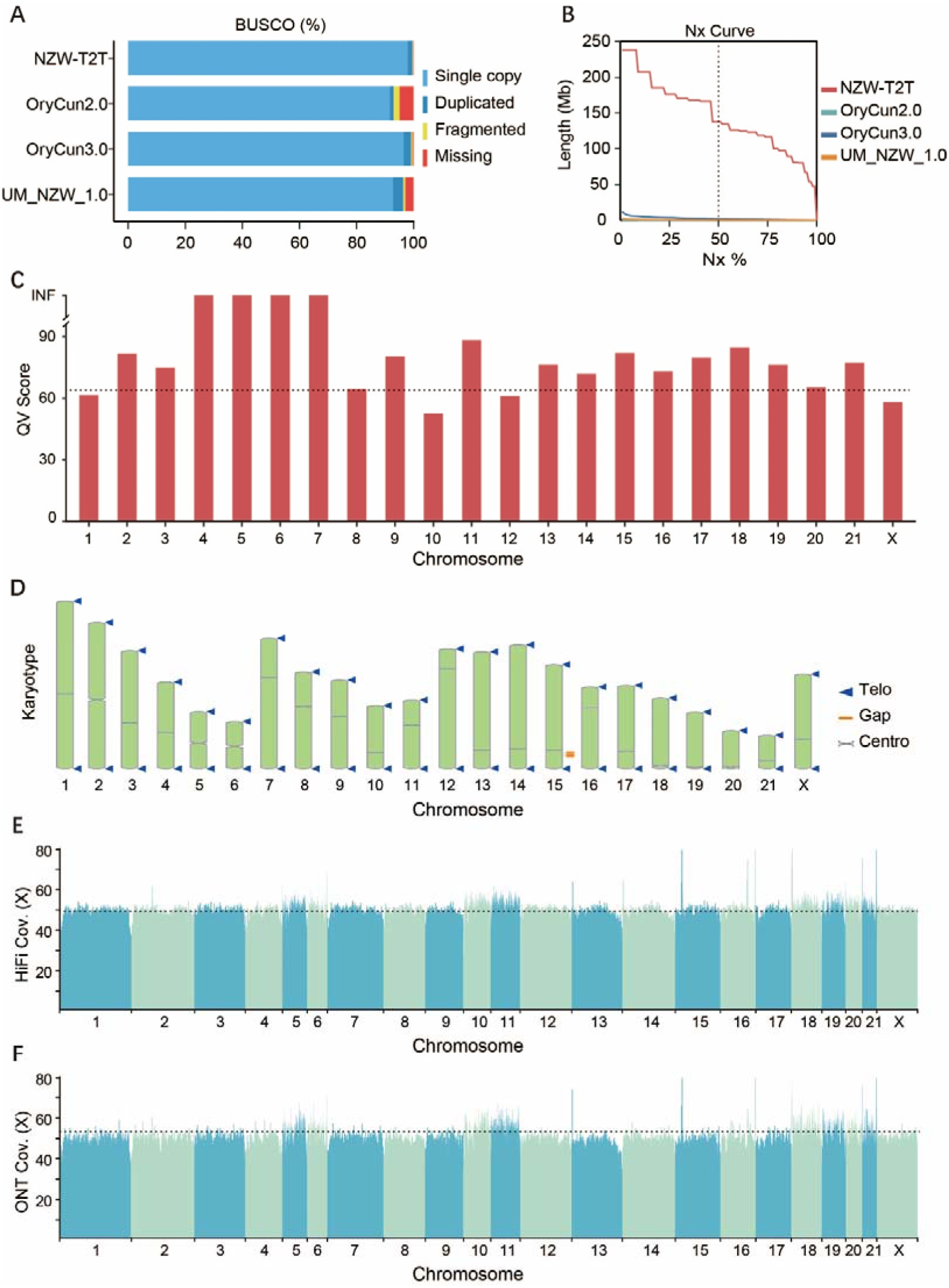
Evaluating the Quality of Genome Assembly. (A) BUSCO completeness assessment results among several rabbit genomes. (B) The assembly contiguity analysis comparison between the NZW-T2T and previous genome assemblies. (C) The QV score, with higher values indicating better quality. The grey line represents the average QV score of the genome. (D) Karyotype representation of the NZW-T2T genome. (E) HiFi sequencing data mapping results. (F) ONT sequencing data mapping results.

In terms of continuity, NZW-T2T exhibited an N50 value of 137.71 Mb, far exceeding OryCun2.0 (64.65 kb), OryCun3.0 (1.88 Mb), and UM_NZW_1.0 (337.73 kb) by 2130.21-fold, 73.29-fold, and 407.76-fold, respectively (Fig. 2B). Quality value (QV) based on k-mer database calculated using Merqury (Rhie et al., 2020), provided further validation of our NZW-T2T assembly, with a whole-genome QV of 64.01 and infinite QV scores for chromosomes 4, 5, 6, and 7, indicating exceptionally high accuracy (Fig. 2C). Additionally, we identified all 22 centromeric regions and 43 telomeric areas, with only one telomeric sequence missing on Chr20 (Fig. 2D). Furthermore, mapping HiFi and ONT reads to NZW-T2T demonstrated a very high mapping rates of 99.99% and 99.18%, respectively, and with an overall even reads depth across the whole genome (Fig. 2E, Fig2F) Overall, to the best of our knowledge, the NZW-T2T assembly represents the most accurate and complete rabbit genome to date.

### Genome annotations

We then performed de novo annotation for the newly assembled genome at both the genomic structure and transcriptome levels. For the genomic features annotations, we applied ab initio prediction and eukaryotic repeat database-based annotation, and identified 1.3 Gb of repetitive sequences, representing 47.40% of the total genome length. The four major types of transposable elements (TEs) detected in the NZW-T2T assembly included short interspersed nuclear elements (SINEs, 19.26%), long interspersed nuclear elements (LINEs, 16.89%), long terminal repeats (LTRs, 5.08%), and DNA transposons (2.08%) (Fig. 3A). Compared to UM_NZW_1.0, OryCun2.0, and OryCun3.0, the genome exhibits proportionally similar contents of the four major transposable element classes (Fig. 3A). In comparison with the UM_NZW_1.0 reference genome –the most complete rabbit genome assembly so far, we additionally identified 185.84 Mb of novel regions in our NZW-T2T assembly, 68.11% of which consisted of repetitive elements. This observation suggests that the complexity of repetitive sequences poses significant challenges for sequencing and assembling.

**Fig. 3.**
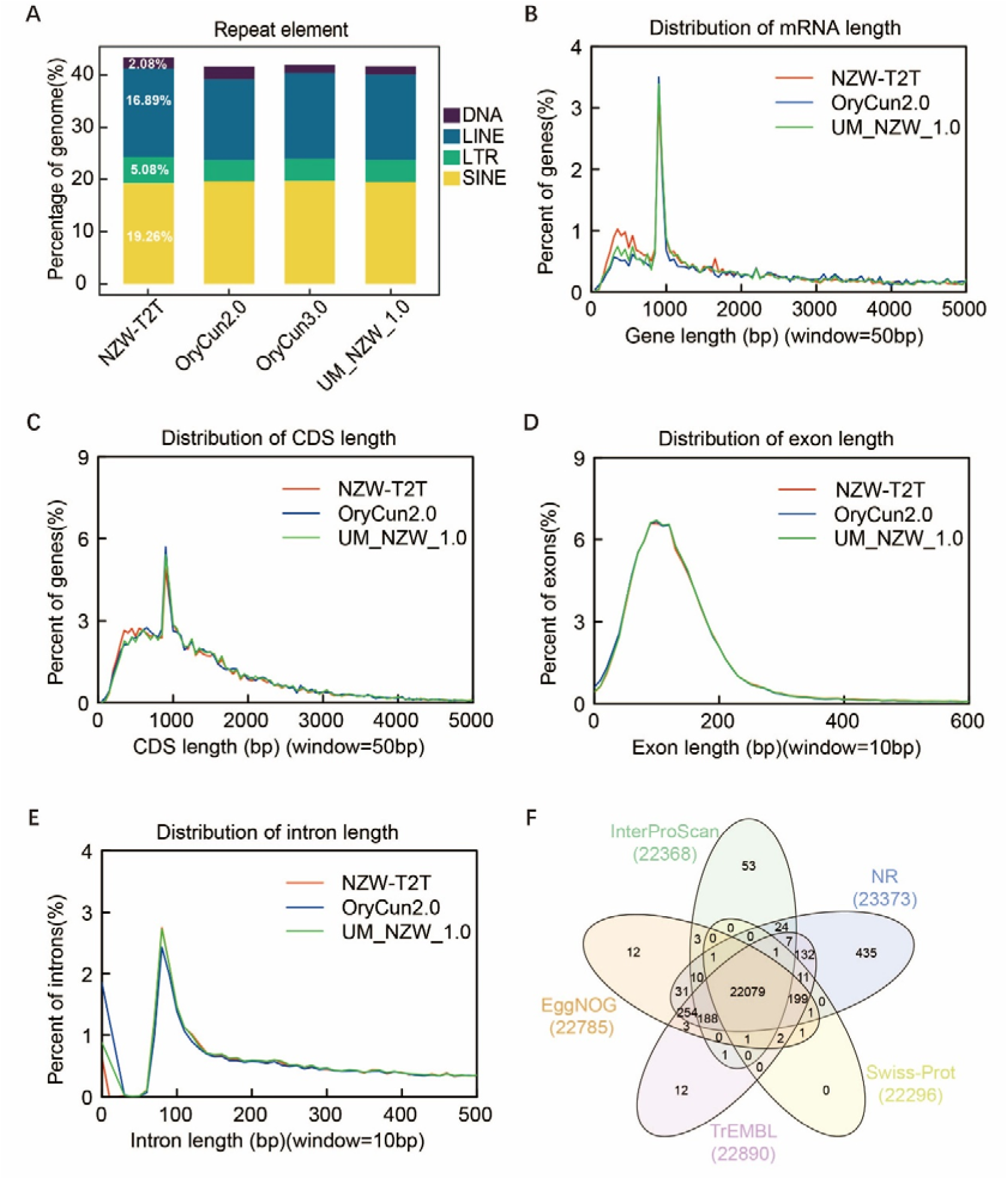
Genome Annotation Overview. (A) Annotation of repetitive sequences across different assemblies and their proportions relative to the total genome size. The proportion of major repeats in the NZW-T2T assembly is marked on the figure. (B-E) Distribution of mRNA, CDS, exon, and intron lengths in the NZW-T2T, UM_NZW_1.0 and OryCun2.0 assembly. (F) Gene function annotation based on five protein databases.

Combined with RNA-Seq data from 7 tissues (Table S8) in this study, we did comprehensive gene annotations. We predicted 23,906 protein-coding genes with an average gene body length of 40,304.1 bp. The distribution of gene lengths and gene content percentages in NZW-T2T assembly were comparable to those in OryCun2.0 and UM_NZW_1.0 (Fig. 3B-E). However, NZW-T2T showed a slightly higher proportion of genes shorter than 1 kb compared to the other two assemblies (Fig. 3B). Predicted genes in NZW-T2T contained an average of 8.5 exons, with mean exon and intron lengths of 184.13 bp and 5,142.56 bp, respectively (Fig. 3D, E). The lengths of gene components, including coding sequences (CDS), exons, and introns, were highly consistent among NZW-T2T, OryCun2.0, and UM_NZW_1.0 (Fig. 3B-E). Functional annotation of the protein-coding genes was conducted by aligning protein sequences against five public databases, including the NCBI non-redundant (NR) protein database, InterPro, SwissProt, TrEMBL, and EggNOG. Of the predicted genes, 22079 (92.36%) were successfully annotated with homologous matches across all five databases; 23,461 (98.14%) were successfully annotated with at least one homologous match across these databases (Fig. 3F).

### NZW-T2T is the state-of-the-art genome assembly of rabbit

The complete, telomere-to-telomere genome assembly holds an extraordinary trove for understanding the development, physiology, and evolution of a species. To evaluate the improvements achieved in the newly assembled genome sequences, we performed a pairwise alignment between our NZW-T2T assembly and UM_NZW_1.0, the most recently published rabbit genome assembly (Table 1). The UM_NZW_1.0 assembly was previously reported to contain 3,137 unplaced contigs, totaling 562.64 Mb and representing 19.8% of the genome (Bai et al., 2021). In contrast, except for one chromosome, NZW-T2T has successfully assembled nearly all chromosomes, with telomeres present at both ends (Fig. 2D), this completeness resolved the majority (99.97%) of unplaced contigs in the UM_NZW_1.0 assembly (Fig. 4A). For example, on chrX, the new assembly resolved 30 unplaced contigs from the UM_NZW_1.0 assembly and their correctness was confirmed by the Hi-C contact heatmap (Fig. 4B). Inversions were also correctly resolved in our new assembly, a large inversion with 33.84 MB on chromosome X in UM_NZW_1.0 was corrected in NZW-T2T, was then confirmed by the Hi-C contact heatmap (Fig. 4B).

**Table 1.**
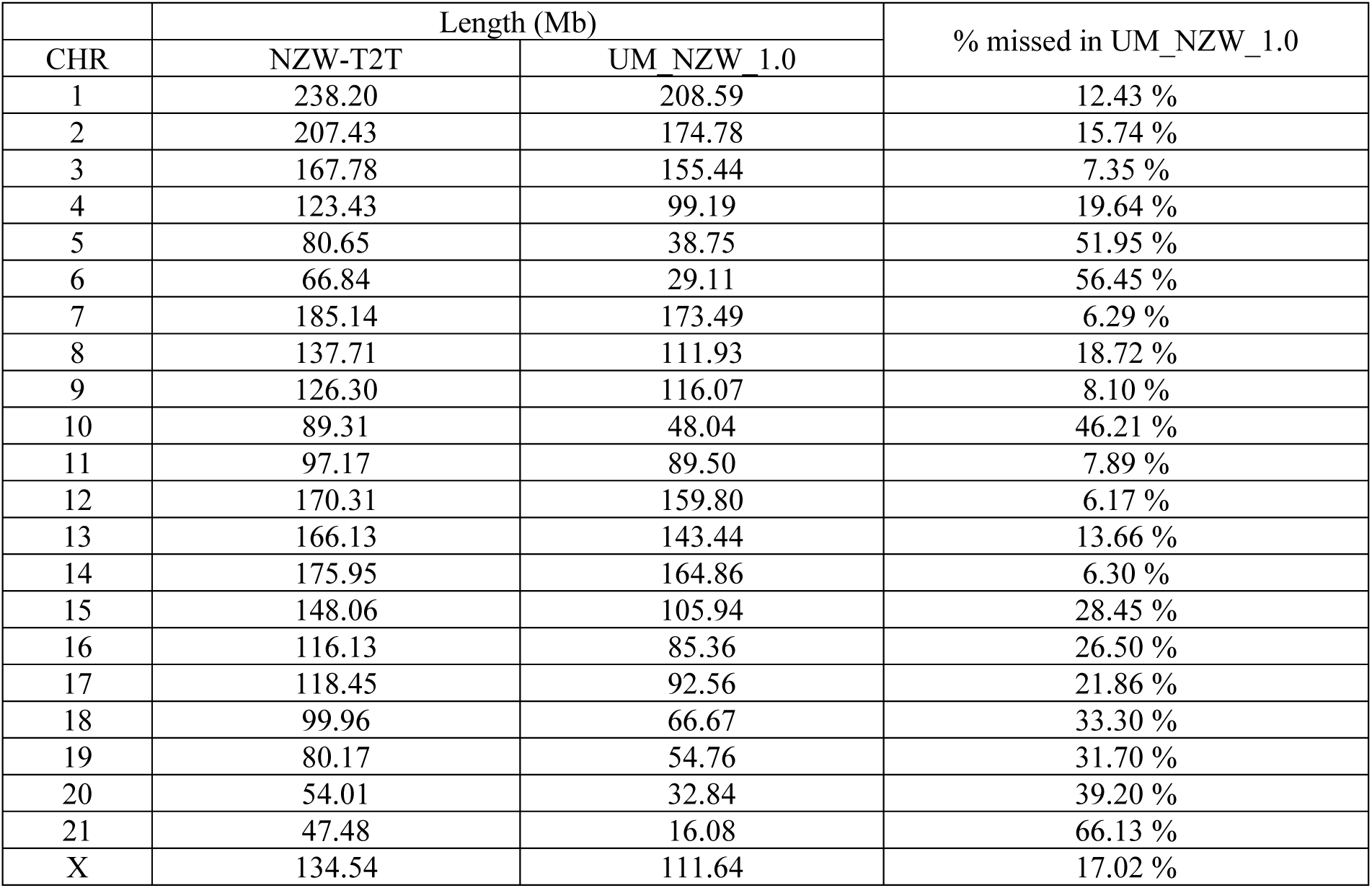
Comparison of chromosome lengths between the NZW-T2T and the UM_NZW_1.0.

| CHR | Length (Mb) |  | % missed in UM_NZW_1.0 |
| --- | --- | --- | --- |
|  | NZW-T2T | UM_NZW_1.0 |  |
| 1 | 238.20 | 208.59 | 12.43 % |
| 2 | 207.43 | 174.78 | 15.74 % |
| 3 | 167.78 | 155.44 | 7.35 % |
| 4 | 123.43 | 99.19 | 19.64 % |
| 5 | 80.65 | 38.75 | 51.95 % |
| 6 | 66.84 | 29.11 | 56.45 % |
| 7 | 185.14 | 173.49 | 6.29 % |
| 8 | 137.71 | 111.93 | 18.72 % |
| 9 | 126.30 | 116.07 | 8.10 % |
| 10 | 89.31 | 48.04 | 46.21 % |
| 11 | 97.17 | 89.50 | 7.89 % |
| 12 | 170.31 | 159.80 | 6.17 % |
| 13 | 166.13 | 143.44 | 13.66 % |
| 14 | 175.95 | 164.86 | 6.30 % |
| 15 | 148.06 | 105.94 | 28.45 % |
| 16 | 116.13 | 85.36 | 26.50 % |
| 17 | 118.45 | 92.56 | 21.86 % |
| 18 | 99.96 | 66.67 | 33.30 % |
| 19 | 80.17 | 54.76 | 31.70 % |
| 20 | 54.01 | 32.84 | 39.20 % |
| 21 | 47.48 | 16.08 | 66.13 % |
| X | 134.54 | 111.64 | 17.02 % |

**Fig. 4.**
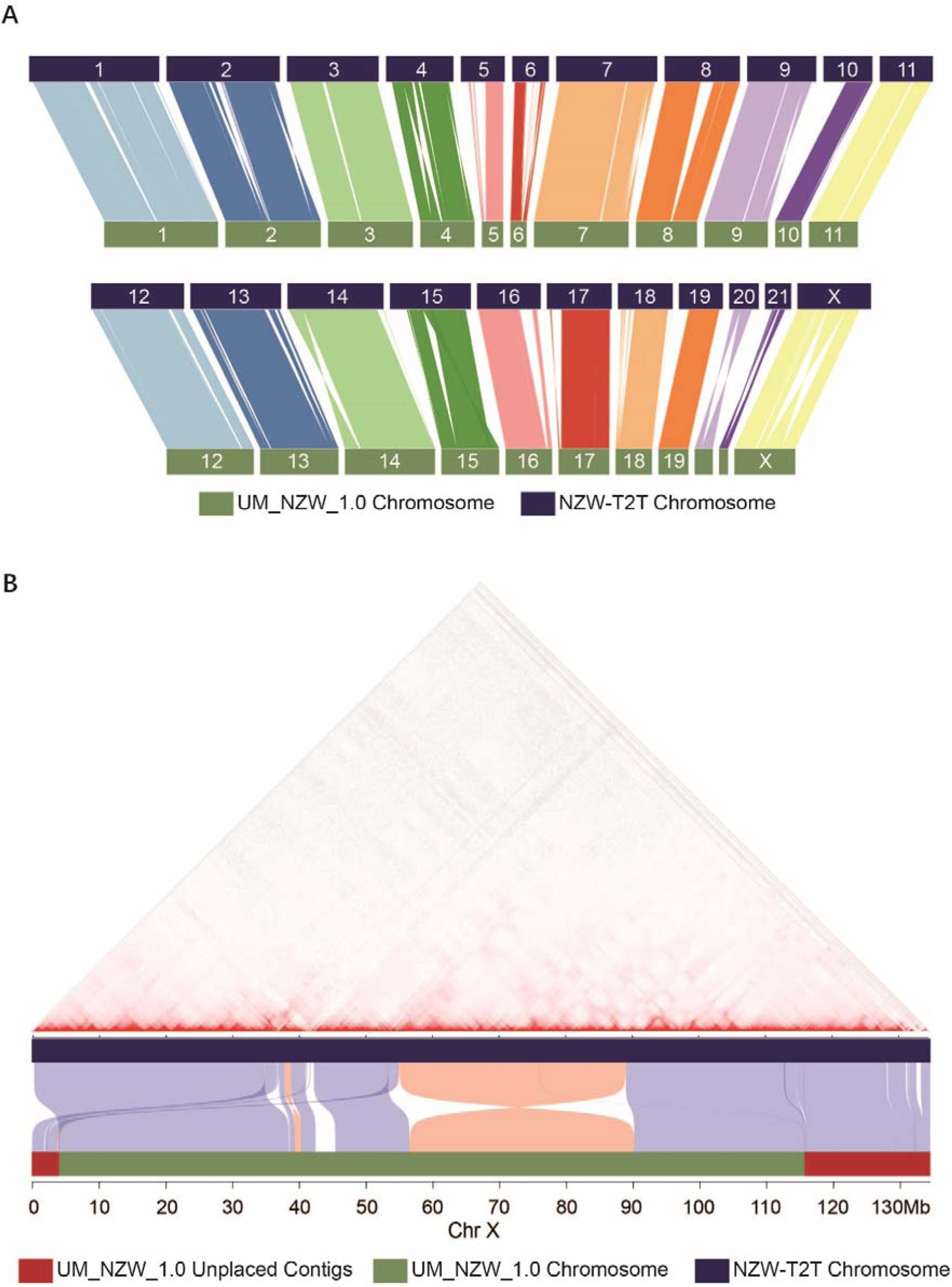
The overview of genomic variations between NZW-T2T and UM_NZW_1.0. (A) Alignments between the chromosomes of NZW-T2T and UM_NZW_1.0. All chromosomes are scaled to their original lengths, with alignments of lengths greater than 100 kb and identities over 99% connected by lines between the chromosomes. (B) Alignment of ChrX between NZW-T2T and UM_NZW_1.0. The upper triangular panel displays a heatmap illustrating Hi-C interactions within ChrX of the NZW-T2T genome. For comparison, the corresponding region in UM_NZW_1.0 is aligned at the bottom of the map. Unplaced contigs in UM_NZW_1.0, UM_NZW_1.0 chromosomes, and NZW-T2T chromosomes are represented in dark red, green, and blue, respectively.

### Fully assembly of immunoglobulin genes

Loci of immunoglobulins represent some of the most complex genomic structures in mammals, particularly the variable genes of the immunoglobulin heavy chain (IGHV). Due to their high sequence similarity and close genomic proximity, no rabbit genome assembly to date has completely and accurately resolved these loci. In the NZW-T2T assembly, the fully resolved genomic structure of all V-D-J-C genes spanning 4.15 Mb in the IgH locus, which is located at one end of chromosome 20 (Fig. 5A). Benefiting from T2T assembly of the Chromosome 20, we successfully identified 367 IGHV genes, 6 IGHD genes, 5 IGHJ genes, and 59 IGHC genes (Fig. 5A). In addition to gene count, but for the first time, all gaps in this region have been completely filled, making it the most complete representation of IgH genes to date.

**Fig. 5.**
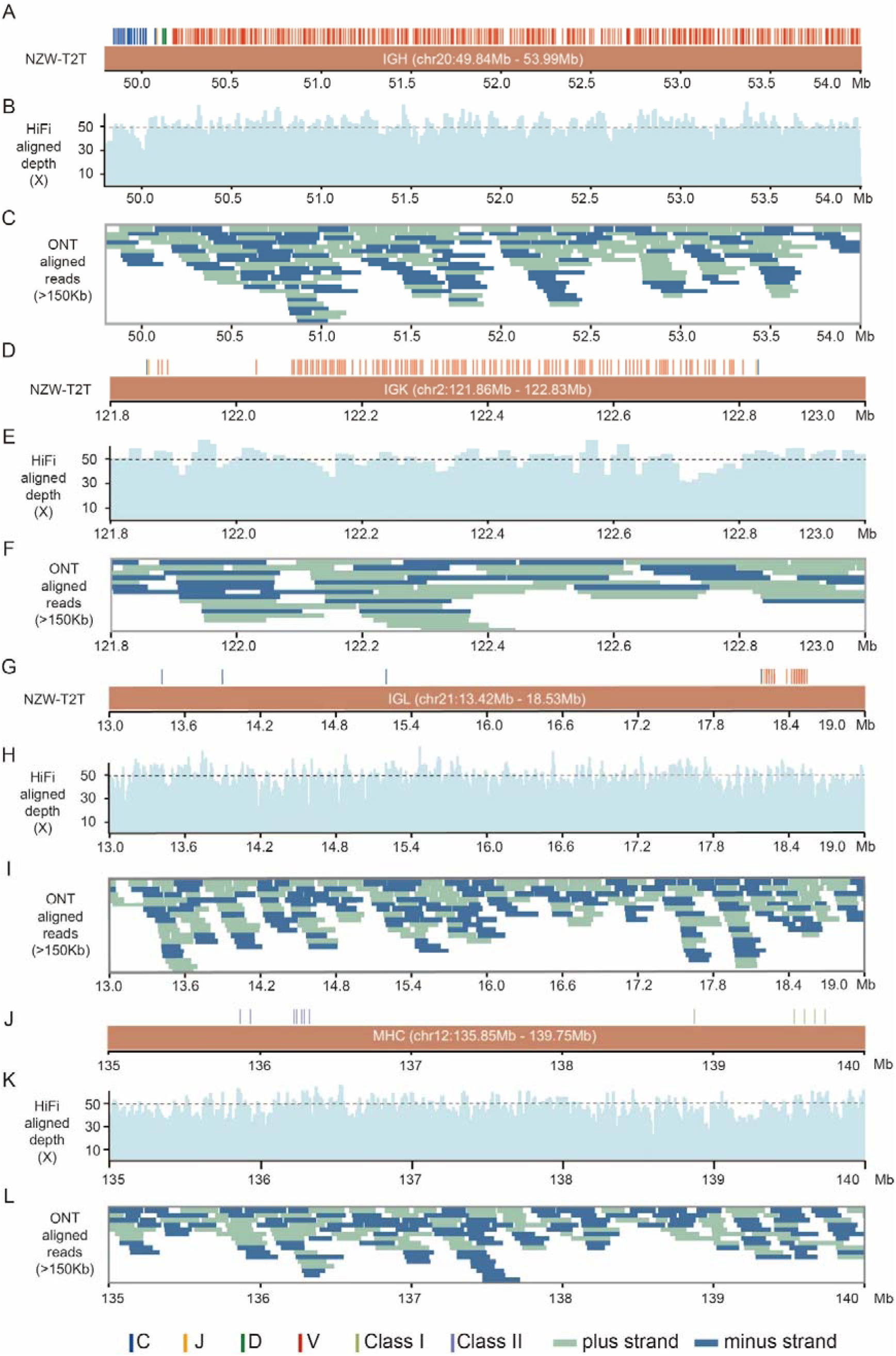
Identification of Ig and MHC genes. The immunoglobulin gene loci are shown for IGH (A), IGK (D), and IGL (G), with V/D/J/C genes represented as red, green, orange, and blue blocks, respectively. (J) shows the MHC region, MHC subregions I and II are marked in light green and purple, respectively. (B), (E), (H)and (K) show the alignment of HiFi reads in the IGH, IGK, IGL and MHC regions, respectively. (C), (F), (I) and (L) show the coverage of Oxford Nanopore ultra-long reads (length > 150 kb) in the IGH, IGK, IGL and MHC regions, respectively.

We confirmed that the two loci for IgK, IgK1 and IgK2, were organized in a head-to-head arrangement on chromosome 2, which had not be described in the previous study (Bai et al., 2021), containing 129 IGKV, 8 IGKJ, and 2 IGKC genes (Fig. 5D). For the IGL genes, which are located on chromosome 21, 46 IGLV, 4 IGLJ, and 6 IGLC genes were identified (Fig. 5G). To assess the accuracy of the assembled immunoglobulin loci, we aligned HiFi reads to the NZW-T2T assembly. Our results showed that read alignments across the IGH, IGK, and IGL gene regions were highly uniform, with HiFi read depths fluctuating within 7.53 standard deviations and ONT read depths within 7.78 standard deviations of the overall sequencing depth (Fig. 5B, Fig. 5C, Fig. 5E, Fig. 5F, Fig. 5H, Fig. 5I).

### MHC gene locus

The major histocompatibility complex (MHC) genes are integral to the adaptive immune response, facilitating antigen presentation and mediating the activation of T cells in response to pathogenic infection. MHC genes primarily consist of two clusters, MHC class I and MHC class II (Petersdorf & O’hUigin, 2019). We located the classical MHC genes in chromosome 12 of NZW-T2T, which aligns with findings from previous studies (Rogel-Gaillard et al., 2001). The rabbit’s classical MHC region revealed by our assembly span 3.9 Mb, while only 3.1 Mb in that of UM_NZW_1.0. The CDS of MHC class I region in rabbits consists of 5 highly polymorphic classical MHC class I genes with a length of 877.46 Kb and while the CDS of MHC class II region contains 7 classical MHC class II genes with a length of 478.10 Kb, (Fig. 5J). Comparative analysis of MHC class I gene sequences from New Zealand rabbits with those of humans, mice, and pigs revealed a sequential decline in homology. Specifically, the homology was highest between humans and rabbits, followed by pigs, and lowest with mice (Fig. S8).

## Discussion

In this study, we successfully generated the first telomere-to-telomere (T2T) genome assembly of the New Zealand White (NZW) rabbit using a haploid embryonic stem cells (haESCs). Compared to previous rabbit genome assemblies, including OryCun1.0, OryCun2.0, and UM_NZW_1.0, the NZW-T2T assembly demonstrates superior completeness, continuity, and accuracy. The near-complete coverage of the BUSCO mammalian gene set (99.61%) highlights the high fidelity of the assembly (Fig. 2A), while the high mapping rates of HiFi (99.99%) and ONT reads (99.18%) further validate its accuracy and completeness (Fig. 2E, Fig. 2F). Additionally, our approach resolves a significant number of previously unanchored contigs, reducing genomic fragmentation and improving chromosomal integrity (Fig. 2D). In the future, the three unresolved gaps on chromosome 15 could be filled by nanopore Cas9-targeted sequencing (nCATS), an enrichment strategy that uses targeted cleavage of chromosomal DNA with Cas9 to ligate adapters for nanopore sequencing (Gilpatrick et al., 2020).

The use of haESCs played a crucial role in enhancing genome assembly accuracy and continuity. In diploid genomes, heterozygosity, segmental duplications, and large structural variations often complicate genome reconstruction (Li & Durbin, 2024). In contrast, haploid cells, which contain only a single set of chromosomes, provide a uniform genetic background (Fig. 1A) (Wang et al., 2024), significantly reducing assembly complexity and facilitating the resolution of repetitive and structurally complex regions, such as centromeres, telomeres, and immune-related gene clusters. The advantage of haploid cells in genome has been demonstrated in human T2T assemblies, where complete hydatidiform mole (CHM) cell lines enabled the reconstruction of entire chromosomes and whole genomes with high accuracy (Logsdon et al., 2021; Miga et al., 2020; Nurk et al., 2022). More recently, haESCs with complete uniform homozygous have been leveraged to generate T2T assemblies in mice (Liu et al., 2024) and in crab-eating macaque (Zhang et al., 2025). Our success in generating a haESC-based T2T genome assembly further supports the concept that the utility of haploid-based sequencing is a valuable approach for high-fidelity genome construction.

A key achievement achieved by using this assembly is the complete annotation of two immune-related genes of the rabbit including immunoglobulin and major histocompatibility complex (MHC) loci (Fig. 5). Immunoglobulin genes, particularly the variable heavy (IGHV) genes, are notoriously difficult to resolve due to their high sequence similarity and complex structural arrangements. Our T2T assembly identifies 367 IGHV genes and providing the most comprehensive view of the rabbit immune gene repertoire to date (Fig. 5A). The resolution of MHC class I and class II regions enhances our understanding of rabbit immunogenetics. Notably, our results indicate that the organization of rabbit MHC genes more closely resembles that of humans than rodents (Fig. S8), reinforcing the utility of rabbits as a model organism for immunological research.

In conclusion, our NZW-T2T assembly provides an unprecedentedly complete rabbit genome, serving as a valuable resource for studies in genetics, immunology, and comparative genomics. The use of haESCs for genome assembly offers a promising strategy for achieving high-contiguity assemblies in other organism with complex genome, facilitating further advancements in the field of genomics. Our success in generating a haESC-based T2T genome assembly in rabbits also underscores its potential as a powerful tool for producing high-quality reference genomes in other species.

## Materials and Methods

### Culturing, library preparing, and sequencing of rabbit haploid embryonic stem cells

rbPhESCs were derived from E4 parthenogenetically developed rabbit blastocysts, with haploid cells enriched and maintained using fluorescence-activated cell sorting (FACS). Prior to library preparation, exponentially expanding rbPhESCs were harvested and sorted for haploid cells. Freshly purified haploid cells were then used to extract high molecular weight DNA following the standard protocol provided by the commercial kit (NucleoBond® HMW DNA, 740160.20). For this study, the following libraries were constructed: paired-end 150 bp libraries for the DNBSEQ-T7 platform, used for the whole genome sequencing, RNASeq, and HiC; SMRTbell libraries for the PacBio Revio platform, used for HiFi sequencing; and ONT Ultra Long libraries for the ONT PromethION platform, used for generating ONT Ultra Long reads.

### Data quality control and genome assembly

HiFi sequencing data were subjected to quality control using SMRT Link V12.0 with default parameters. ONT sequencing data were subjected to quality control using Nanoq with default parameters, and reads containing sequencing adapter sequences and those with a quality score below 7 were removed. All HiFi reads and ONT reads that passed quality control were used for genome assembly with hifiasm-ul (Cheng et al., 2024) using default parameters, resulting in the asm_hia assembly. These quality-controlled HiFi and ONT sequences were further assembled using verkko (Rautiainen et al., 2023), producing the asm_vko assembly. The cleaned Hi-C sequencing data, which were processed using the default parameters of Soapnuke software for quality control, were aligned to the contig assembly using Bowtie2 (Langmead & Salzberg, 2012) to identify uniquely mapped paired-end reads. Quality control of the aligned reads using HiC-Pro (Servant et al., 2015) to removed 407,218,431 invalid pairs, representing 47.42% of the cleaned Hi-C sequencing data. Subsequently, quality controlled Hi-C reads were utilized to generate a scaffolded genome assembly with Juicer (Durand, Robinson, et al., 2016) and 3D-DNA (Dudchenko et al., 2017). The assembly was visually inspected and corrected using JuiceBox (Durand, et al., 2016) based on chromosomal interaction strength. Contigs with no clear interaction relationships were treated as individual scaffolds, while those assigned to the same chromosome were concatenated with 500 ‘N’ bases to construct the final chromosome-level genome assembly. For each chromosome from two assembly versions, we assess the assembly quality based on the completeness of telomere assembly, the presence of gaps, and the QV score (Rhie et al., 2020), thereby selecting the superior version. The Bandage software was utilized for visualizing final sequence graphs (Wick et al., 2015).

### Assembly quality assessment

The completeness of the genome assembly was evaluated using the compleasm software (Huang & Li, 2023) with the mammalia_odb10 database to identify conserved single-copy genes in OryCun2.0 (Carneiro et al., 2014), OryCun3.0 (GCF_009806435.1), UM_NZW_1.0 (Bai et al., 2021), and NZW-T2T. The BUSCO plot was generated using the generate_plot.py script from the BUSCO package (Seppey et al., 2019). The assembly N50 value was calculated using calN50 (https://github.com/lh3/calN50), and the Nx curve was plotted using a custom R script. Assembly quality was quantified with HiFi reads using Merqury software (Rhie et al., 2020). We employed the telomere recognition motif “TTAGGG” to identify telomeres, which is specific to animals (Louzon et al., 2019). The TeloExplorer function of the quarTeT software (Lin et al., 2023)was utilized for telomere identification. To identify centromeres, we used the centromics software (https://github.com/ShuaiNIEgithub/Centromics). The locations of the centromeres and telomeres have been shown on the chromosome with R package Rideogram (Hao et al., 2020). PacBio HiFi and ONT ultralong reads were aligned to the NZW-T2T genome using Inspector software (Y. Chen et al., 2021), which yielded overall alignment rates. For each chromosome, coverage depth plots were computed and visualized using Svhawkeye software (Xiao et al., 2024).

### Gene annotation

RepeatModeler (v2.0.4) (Flynn et al., 2020) was employed to detect repetitive elements using default settings, and then a de novo repeat sequence library was established. Additionally, a custom library was created by integrating data from Dfam 3.7 (Hubley et al., 2016) and RepBase20181026 databases (Bao et al., 2015). To perform homology-based annotation, repetitive elements were masked utilizing RepeatMasker (v4.1.5) (Tarailo-Graovac & Chen, 2009) against the custom library. As for gene annotation, we employed a combination of three methods: ab initio prediction, transcriptome-based annotation, and homology-based annotation. Ab initio prediction was performed using the Augustus(v3.5.0) pipeline (Stanke et al., 2008). For transcriptome-based annotation, RNA-seq data from seven different tissues was first mapped to our assembly using HISAT2 (D. Kim et al., 2015) and transcripts were assembled from the alignment data using StringTie (v2.1.7) (Pertea et al., 2015). Finally, TransDecoder (v5.5.0) was applied to identify candidate coding regions and predict protein-coding gene models based on the assembled transcripts. Protein sequences for the European hare, human, and mouse were retrieved from the RefSeq database and used as reference sequences in Genewise (Birney et al., 2004) for homology-based gene structure prediction.The gene prediction model described above is integrated by EvidenceModeler(v1.1.1) (Haas et al., 2008) and supplemented with the genetic annotation obtained by Liftoff(v1.6.3) (Shumate & Salzberg, 2021) to form the final gene annotation.

### Genome synteny analysis

Given that UM_NZW_1.0 contains 3,137 unplaced contigs, we first visualized the 22 chromosomes of UM_NZW_1.0 in comparison with the 22 chromosomes of NZW-T2T using NgenomeSyn (He et al., 2023) with default parameters to highlight the similarities and differences between the two genomes. To address the unplaced contigs from UM_NZW_1.0, we used minimap2 (Li, 2018) to align them to the NZW-T2T reference genome. For each unplaced contig, we determined its most likely chromosomal location based on the highest sequence similarity score, obtaining specific coordinates. Finally, we visualized the synteny relationships between the UM_NZW_1.0 and NZW-T2T genomes using the R package Asynt (K.-W. Kim et al., 2022).

### Immunoglobulin gene mapping

Rabbit nucleotide sequences of the V, J and D genes of IGH/IGK/IGL were obtained from “F + ORF+all P” sets in the IMGT/GENE-DB reference directory, and the nucleotide sequences of the C genes were fetched from the IMGT/GENE-DB query page (Giudicelli et al., 2005). The sequences were aligned to NZW-T2T with BLAST, of which D genes were mapped using “-task blastn-short” due to their short sequence length. For each D, J and C gene, only the BLAST result with the highest score was kept. As for V genes, all results with alignment coverage greater than 0.9 were preserved. In the case of overlapping target regions, the blast result with the smallest e-value was chosen.

### MHC region identification

The human MHC region was aligned to the NZW-T2T genome using minimap2 (v2.22-r1101) (Li, 2018), confirming that the MHC is located on chromosome 12 of the rabbit. The homologous sequences of human, mouse, and rabbit were downloaded from the RefSeq, Ensembl, and Swissprot databases and utilized for the prediction of potential MHC gene loci using genblasta (v1.0.1)(She et al., 2009) with parameter of “-e 1e-5 -c 0.8”, retaining only regions with an alignment coverage of no less than 80%. GeneWise (v2.4.1)(Birney et al., 2004) was then used to re-annotate the aforementioned regions. For homology comparison, we extracted major histocompatibility complex (MHC) class I gene sequences from reference genome annotations in human (T2T-CHM13v2.0), mouse (GRCm39), pig (Sscrofa11.1) and aligned with NZW-T2T using ClustalW (Thompson et al., 2002) The phylogenetic tree of MHC I genes was visualized using iTOL (Letunic & Bork, 2021). Considering the absence of a standardized nomenclature for rabbit MHC class I genes (Pinheiro et al., 2016), we assigned names based on their relative chromosomal positions within the MHC I genomic regions to enhance clarity in presentation.

## Code availability

The code for this study has been uploaded to https://github.com/ayungaa/rabbit-t2t-genome.

## Data availability

The sequencing data generated in this study have been deposited in the Genome Sequence Archive database and are available under the project accession number PRJCA036842.

## Acknowledgements

We thank Junwei Wang, Minghui Gao for the help with FACS. This work was supported by the National Natural Science Foundation of China (32100409, 82100482, 82101553 and 82200504, 82470352), the Science and Technology Planning Project of Guangdong Province (2022A1515012058), the Science and Technology Program of Guangzhou (202201010617), the Funding by Science and Technology Projects in Guangzhou (2024A04J4399), the Noncommunicable Chronic Diseases-National Science and Technology Major Project-2023ZD0503201, Pioneering Action Grants of the Chinese Academy of Sciences, Zhejiang Provincial Clinical Research Center for Pediatric Diseases-ZJEK2302Z.

## Notes

### Competing Interest Statement

The authors have declared no competing interest.

