## Supplemental File for "Construction of Complete Telomere-to-Telomere Genome Assembly of the Rabbit Using Haploid Embryonic Stem Cells"

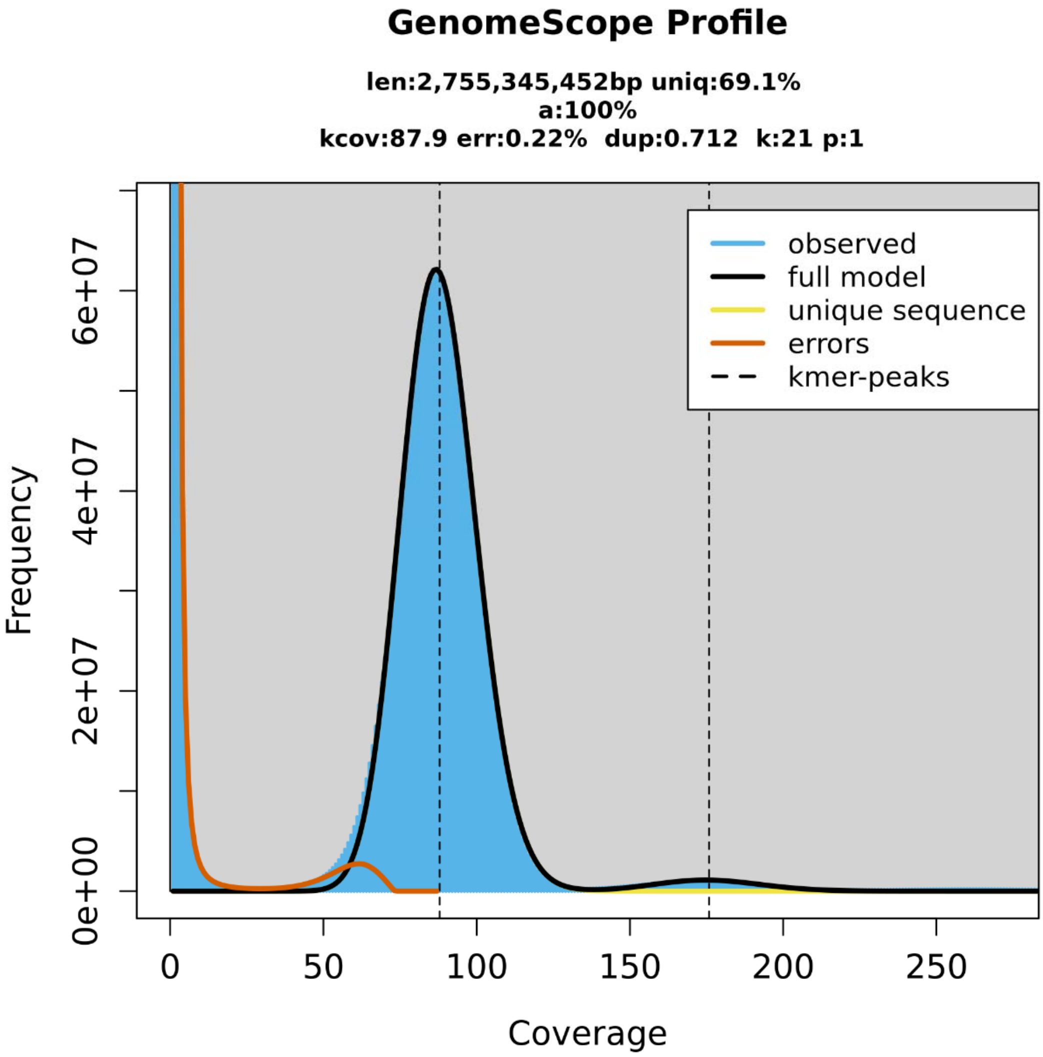


**Supplementary Fig. 1. Genome size estimation**

The blue area represents the actual k-mer distribution. Low-frequency k-mer below the red line are considered sequencing errors, while reliable data below the black line are used for genome size estimation. Vertical dashed lines indicate k-mer peaks, and the area below the yellow line represents the size of non-repetitive regions.

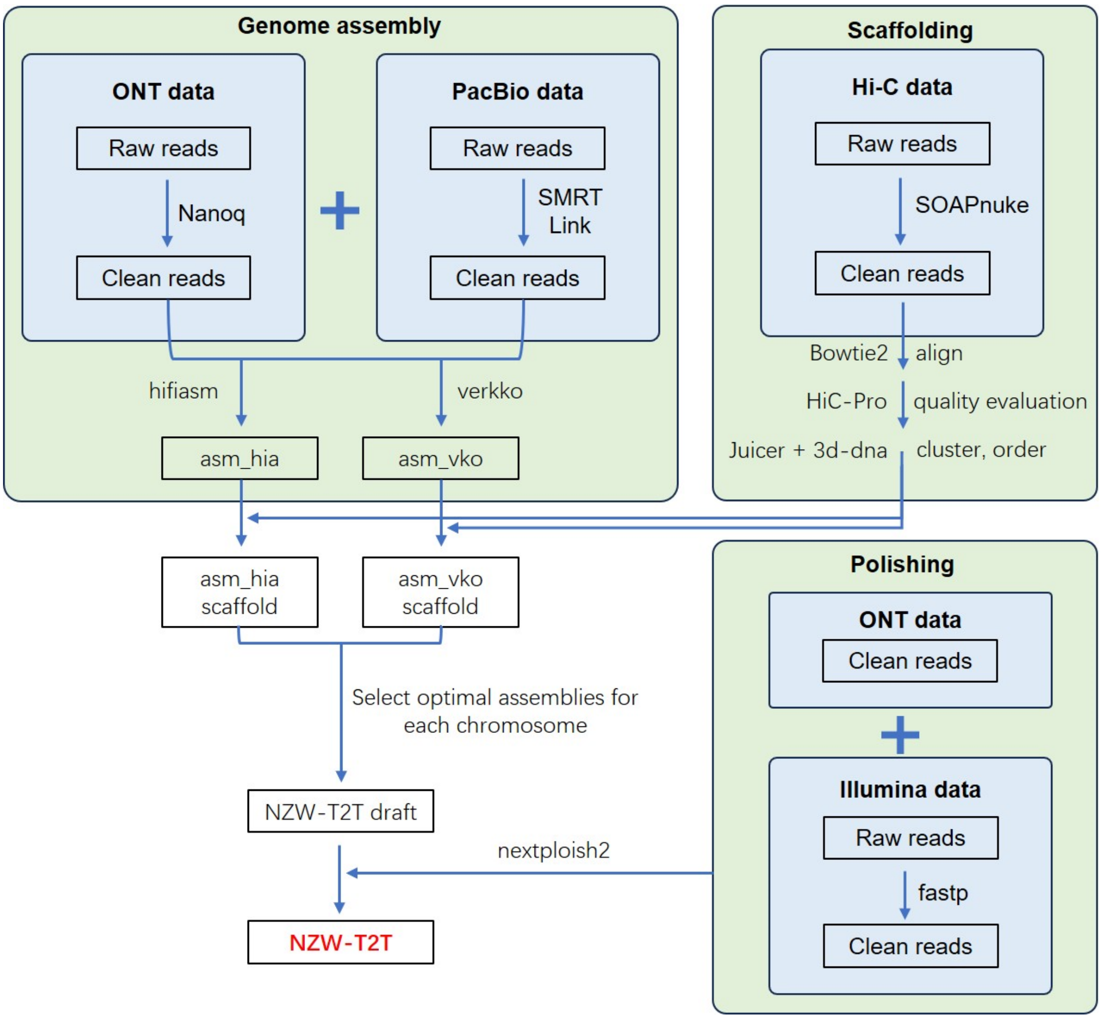


**Supplementary Fig. 2. Genome assembly workflow diagram**

During the construction of the NZW-T2T genome, we first employed two distinct assembly algorithms—hifiasm and verkko—for preliminary genome assembly. Subsequently, for the contigs generated from each preliminary assembly, we performed chromosomal anchoring to obtain the corresponding scaffold sequences. At this point, for each chromosome, we had scaffolds constructed from both assembly algorithms. For each chromosome, we selected the scaffold with the highest assembly quality to represent that chromosome. Finally, we conducted gap filling and polishing analyses to further optimize the quality of the genome assembly.

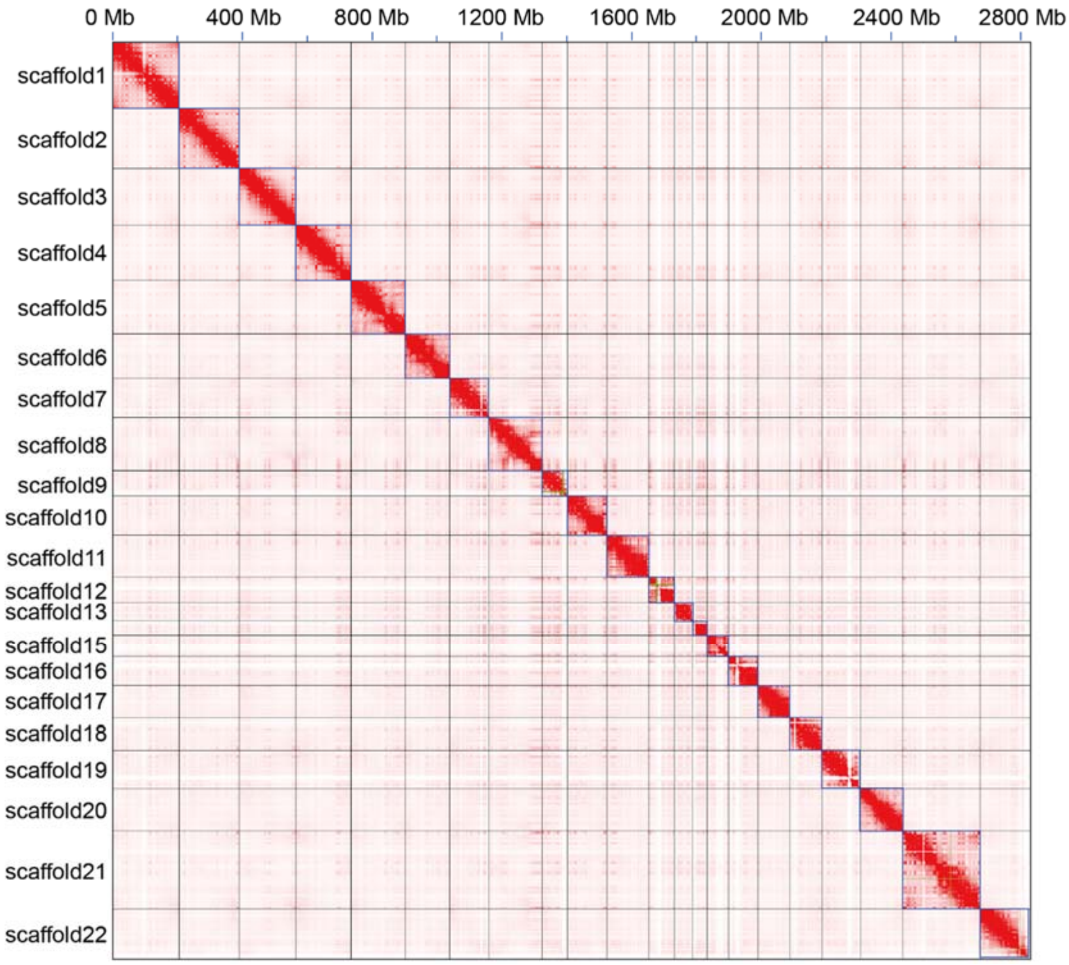


**Supplementary Fig. 3. Chromosome Interaction Map of asm_hia**

The color intensity of each cell in the figure represents the strength of interaction between different genomic regions, with darker colors indicating stronger interactions. Both the x-axis and y-axis represent the scaffolds of asm_hia, arranged sequentially.

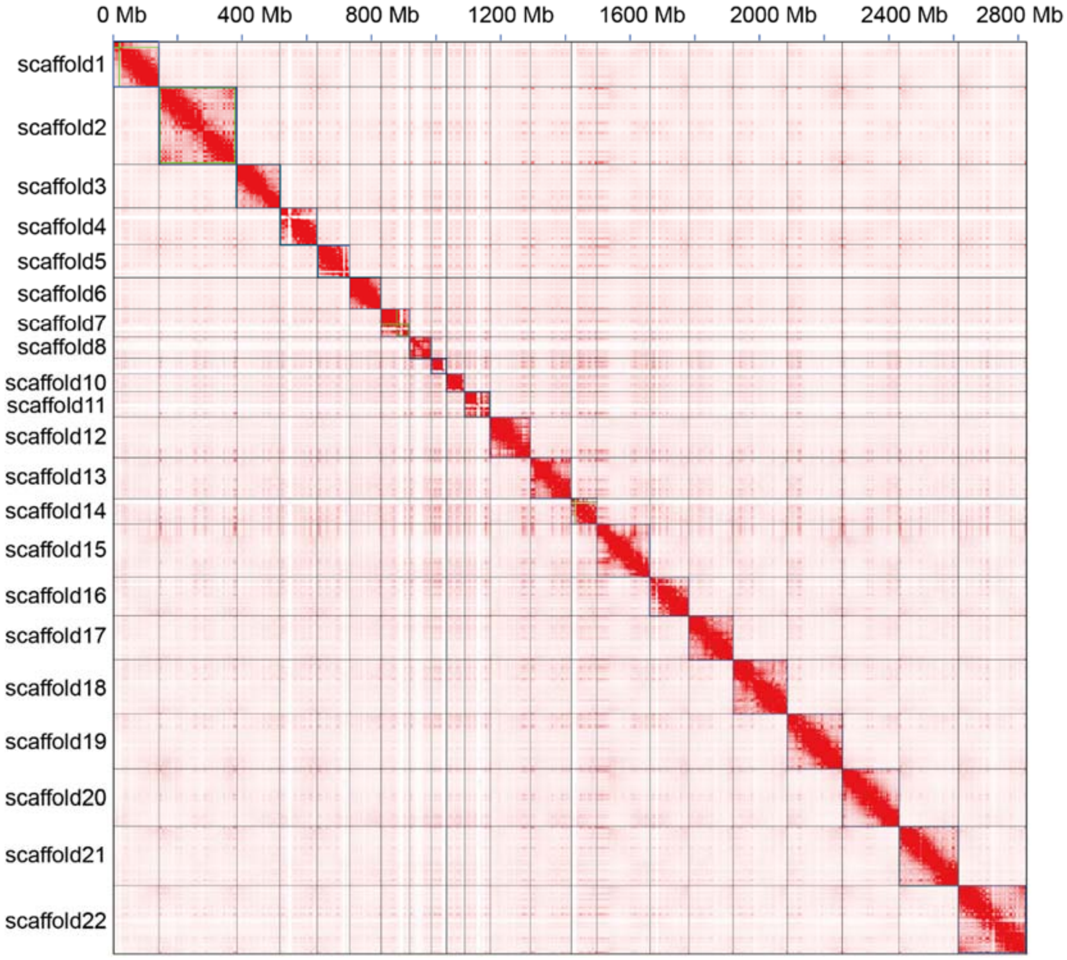


**Supplementary Fig. 4. Chromosome Interaction Map of asm_vko**

The color intensity of each cell in the figure represents the strength of interaction between different genomic regions, with darker colors indicating stronger interactions. Both the x-axis and y-axis represent the scaffolds of asm_vko, arranged sequentially.

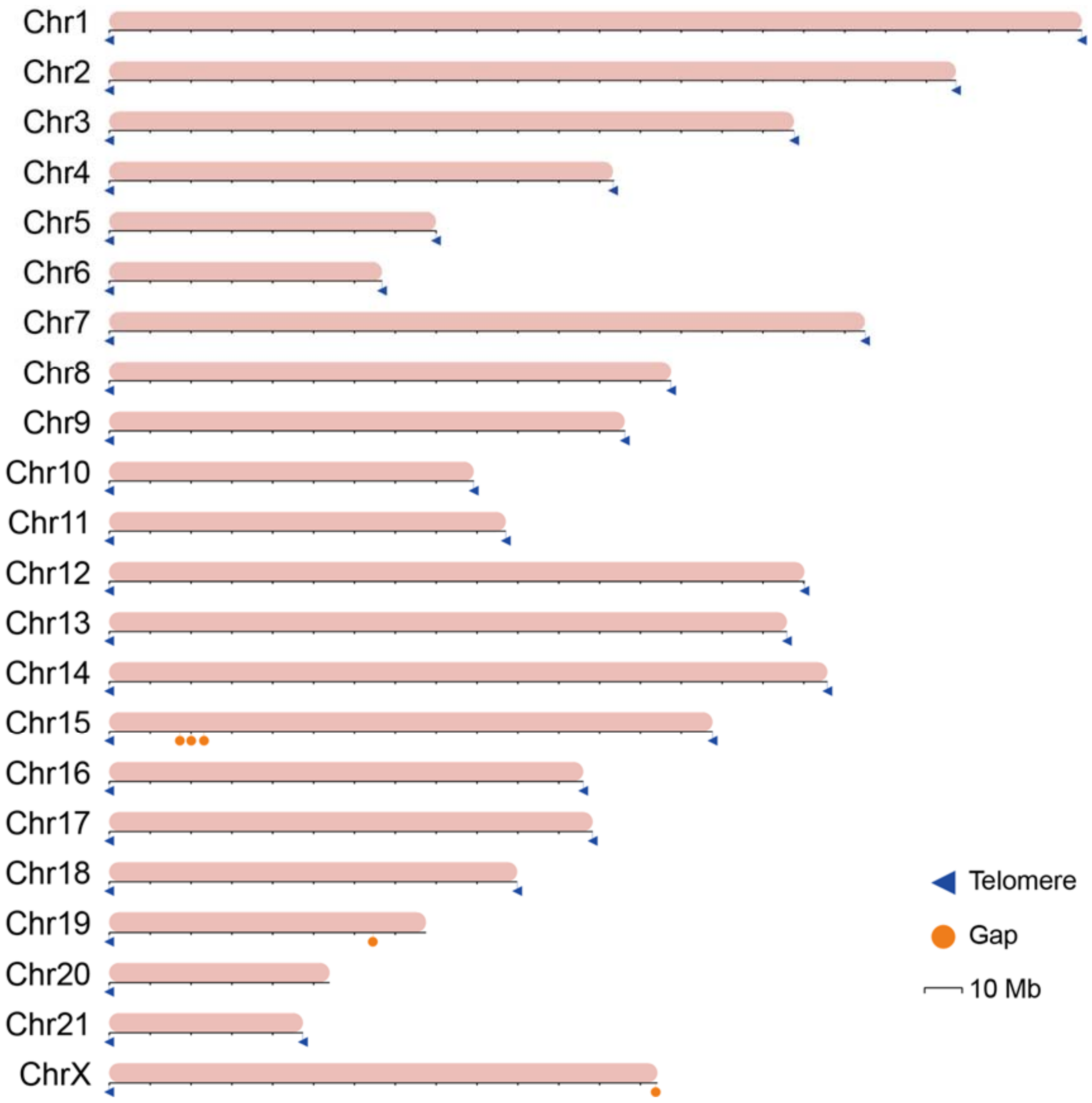


**Supplementary Fig. 5. Schematic diagram of the chromosome of asm_hia**

Each horizontal bar in the figure represents an individual chromosome of asm_hia, with the unit of chromosome length indicated below the bar. Blue triangles are used to mark the positions of telomeres, while yellow circles denote gap regions on the chromosomes.

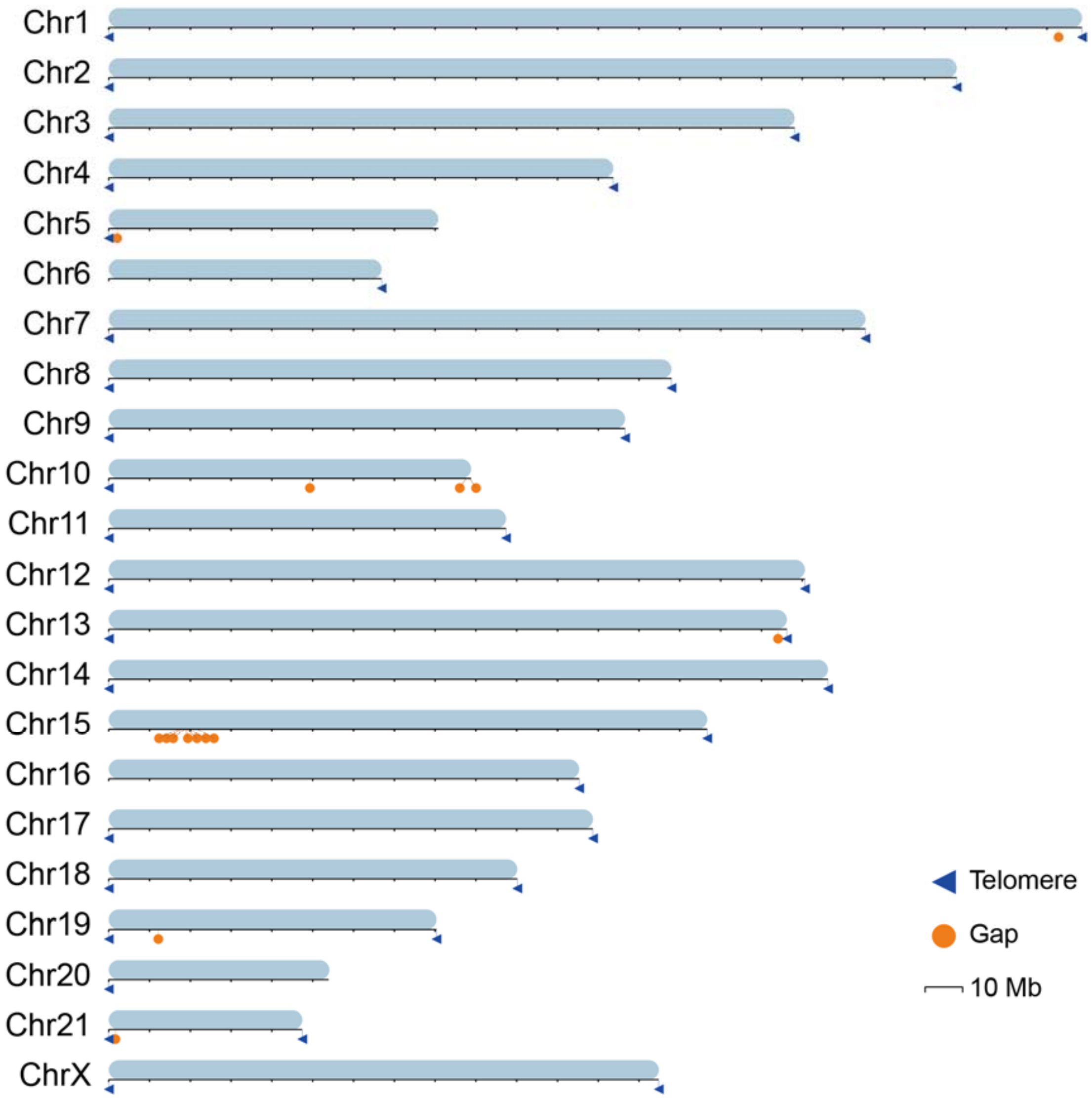


**Supplementary Fig. 6. Schematic diagram of the chromosome of asm_vko**

Each horizontal bar in the figure represents an individual chromosome of asm_vko, with the unit of chromosome length indicated below the bar. Blue triangles are used to mark the positions of telomeres, while yellow circles denote gap regions on the chromosomes.

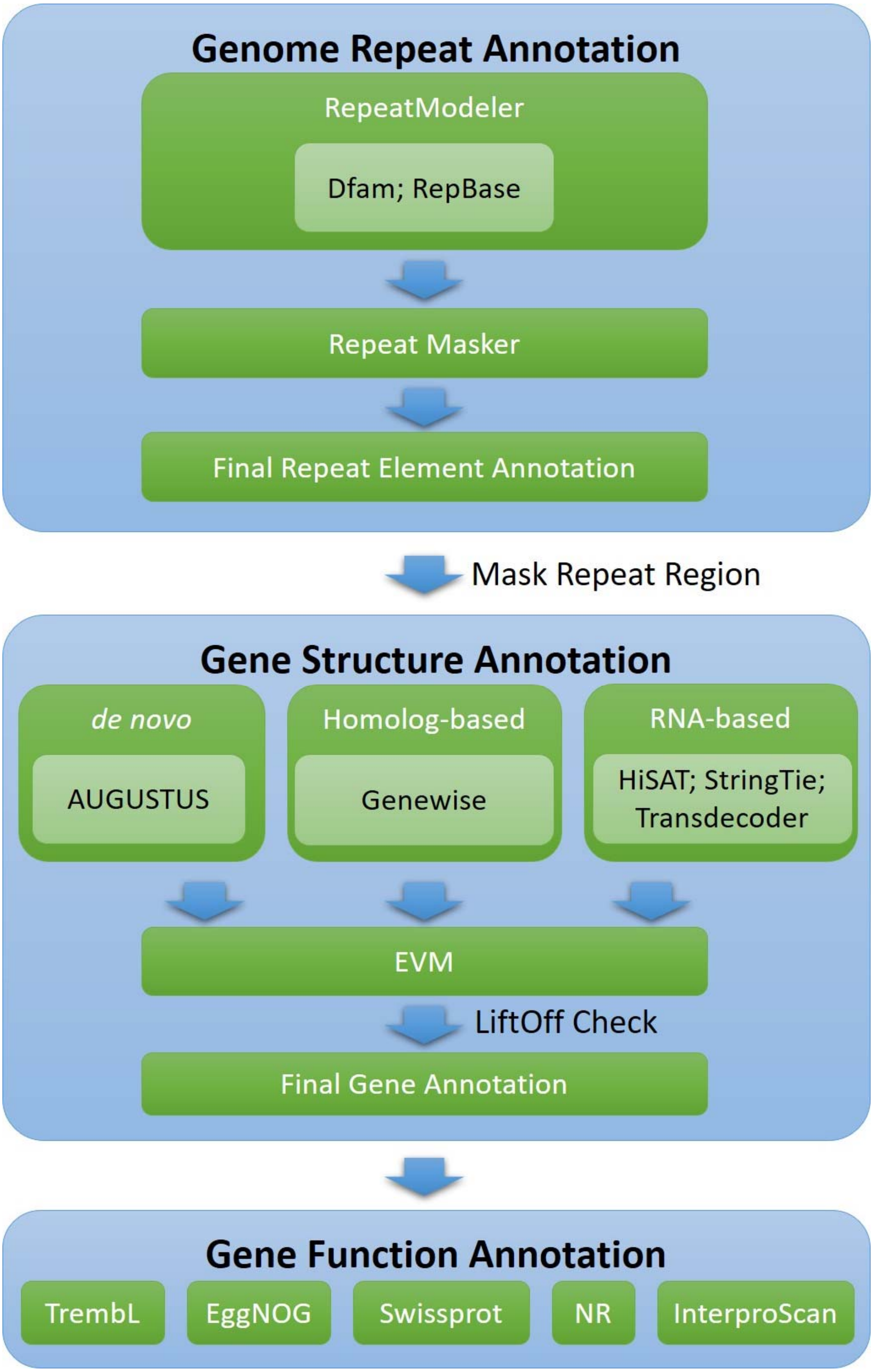


**Supplementary Fig. 7. Genome annotation workflow diagram**

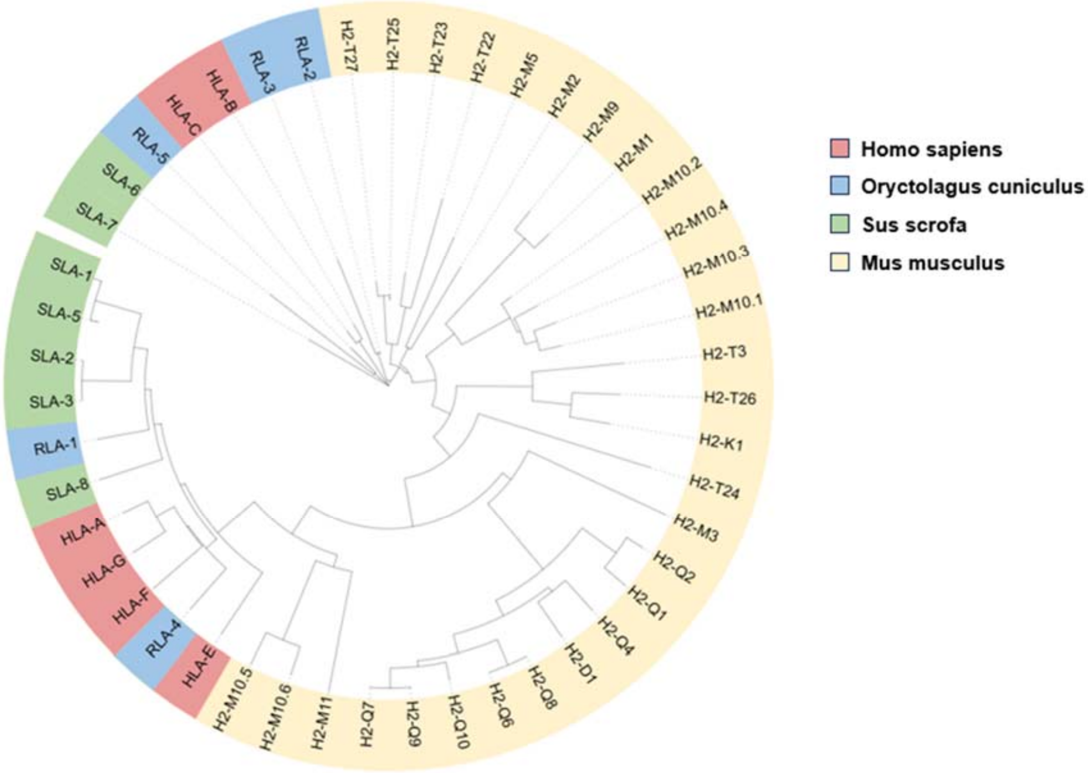


**Supplementary Fig. 8. Phylogenetic tree of MHC class I genes**

**Supplementary Table 1 Quality information for ONT sequencing data**

| Item | Data |
| --- | --- |
| Mean read length | 36.13 kb |
| Mean read quality | 13.8 |
| Median read length | 21.23 kb |
| Median read quality | 14.7 |
| Number of reads | 4,164,493 |
| Read length N50 | 72.93 kb |
| Total bases | 150.49 Gb |
| Number of reads above Q10 | 3,911,754 |
| Percentage of reads above Q10 | 93.9% |
| Size of reads above Q10 | 138.63 Gb |

**Supplementary Table 2 Statistics of read1 quality control for Hi-C sequencing data**

| Item | Read1 (raw) | Read1 (clean) |
| --- | --- | --- |
| Total reads | 858763581 | 858763039 |
| Total bases | 128,814,537,150 | 128,814,455,850 |
| Q20 bases | 126,868,481,390 (98.49%) | 126,868,406,267 (98.49%) |
| Q30 bases | 122,421,007,350 (95.04%) | 122,420,937,958 (95.04%) |

**Supplementary Table 3 Statistics of read2 quality control for Hi-C sequencing data**

| Item | Read1 (raw) | Read2 (clean) |
| --- | --- | --- |
| Total reads | 858,763,581 | 858,763,039 |
| Total bases | 128,814,537,150 | 128,814,455,850 |
| Q20 bases | 125,479,228,478 (97.41%) | 125,479,162,539 (97.41%) |
| Q30 bases | 118,227,354,039 (91.78%) | 118,227,299,635 (91.78%) |

**Supplementary Table 4 Statistics of read1 quality control for DNA next-generation sequencing data**

| Item | Read1 (raw) | Read1 (clean) |
| --- | --- | --- |
| Total reads | 440,568,211 | 440,564,842 |
| Total bases | 66,085,231,650 | 65,901,775,414 |
| Q20 bases | 65,124,166,924 (98.5457%) | 64,943,498,454 (98.5459%) |
| Q30 bases | 62,843,375,856 (95.0944%) | 62,669,240,237 (95.0949%) |

**Supplementary Table 5 Statistics of read2 quality control for DNA next-generation sequencing data**

| Item | Read2 (raw) | Read2 (clean) |
|---|---|---|
| Total reads | 440,568,211 | 440,564,842 |
| Total bases | 66,085,231,650 | 65,901,775,414 |
| Q20 bases | 63,937,539,314 (96.7501%) | 63,762,113,440 (96.7533%) |
| Q30 bases | 59,456,455,394 (89.9694%) | 59,294,380,811 (89.9739%) |

**Supplementary Table 6 Statistics of contig number and telomere number of each chromosome for asm_hia and asm_vko**

| Chromosome ID | Number of contigs in asm_hia (#) | Number of telomere in asm_hia (#) | Number of contigs in asm_vko (#) | Number of telomere in asm_hia (#) |
|---|---|---|---|---|
| Chr 1 | 1 | 2 | 2 | 2 |
| Chr 2 | 1 | 2 | 1 | 2 |
| Chr 3 | 1 | 2 | 1 | 2 |
| Chr 4 | 1 | 2 | 1 | 2 |
| Chr 5 | 1 | 2 | 2 | 1 |
| Chr 6 | 1 | 2 | 1 | 1 |
| Chr 7 | 1 | 2 | 1 | 2 |
| Chr 8 | 1 | 2 | 1 | 2 |
| Chr 9 | 1 | 2 | 1 | 2 |
| Chr 10 | 1 | 2 | 4 | 1 |
| Chr 11 | 1 | 2 | 1 | 2 |
| Chr 12 | 1 | 2 | 1 | 2 |
| Chr 13 | 1 | 2 | 2 | 2 |
| Chr 14 | 1 | 2 | 1 | 2 |
| Chr 15 | 4 | 2 | 8 | 1 |
| Chr 16 | 1 | 2 | 1 | 1 |
| Chr 17 | 1 | 2 | 1 | 2 |
| Chr 18 | 1 | 2 | 1 | 2 |
| Chr 19 | 2 | 1 | 2 | 2 |
| Chr 20 | 1 | 1 | 1 | 1 |
| Chr 21 | 1 | 2 | 2 | 2 |
| Chr X | 2 | 1 | 1 | 2 |

**Supplementary Table 7 Four types of transposable elements located in the new region**

| Repeat Type | Length (Mb) | Percent (%) |
|---|---|---|
| DNA | 1.47 | 0.79 |
| LINE | 39.66 | 21.34 |
| SINE | 27.27 | 14.67 |
| LTR | 8.58 | 4.62 |

**Supplementary Table 8 Statistics of sequencing data volume for RNA-seq from different tissues**

| Sample tissue name | Sequencing type | Clean data |
|---|---|---|
| Thymus gland | RNA-seq | 14.19 Gb |
| Gluteus maximus | RNA-seq | 17.91 Gb |
| Adrenal gland | RNA-seq | 15.72 Gb |
| Ovary | RNA-seq | 13.87 Gb |
| Cerebral hemisphere | RNA-seq | 16.88 Gb |
| Colon | RNA-seq | 15.26 Gb |
| Testis | RNA-seq | 15.01 Gb |
